# Pooled genome-wide CRISPR screening for basal and context-specific fitness gene essentiality in *Drosophila cells*

**DOI:** 10.1101/274464

**Authors:** Raghuvir Viswanatha, Zhongchi Li, Yanhui Hu, Norbert Perrimon

## Abstract

Genome-wide screens in *Drosophila* cells have offered numerous insights into gene function, yet a major limitation has been the inability to stably deliver large multiplexed DNA libraries to cultured cells allowing barcoded pooled screens. Here, we developed a site-specific integration strategy for library delivery and performed a genome-wide CRISPR knockout screen in *Drosophila* S2R+ cells. Under basal growth conditions, 1235 genes were essential for cell fitness at a false-discovery rate of 5%, representing the highest-resolution fitness gene set yet assembled for *Drosophila*, including 407 genes which likely duplicated along the vertebrate lineage and whose orthologs were underrepresented in human CRISPR screens. We additionally performed context-specific fitness screens for resistance to or synergy with trametinib, a Ras/ERK/ETS inhibitor, or rapamycin, an mTOR inhibitor, and identified key regulators of each pathway. The results present a novel, scalable, and versatile platform for functional genomic screens in low-redundancy animal cells.

## Introduction

Systematic perturbation of gene function in eukaryotic cells using arrayed (well-by-well) reagents is a powerful technique that has been used to successfully assay many fundamental biological questions such as proliferation, protein secretion, morphology, organelle maintenance, viral entry, synthetic lethality, and other topics (1). An alternative approach, widely used in mammalian cells, is pooled screening that uses limited titers of integrating lentiviral vectors carrying a perturbative DNA sequence such that each cell receives one integrating virus. In pooled screens, the perturbing DNA reagent serves as the tag in subsequent sequencing (2-4). A key benefit of this approach is that pool size can be extremely large, allowing high reagent multiplicity and thus increased screen quality. The pooled approach in mammalian cells has been used extensively to perform RNAi and more recently single guide RNA (sgRNA) screens using CRISPR/Cas9 (5-7).

Genetic loss-of-function arrayed RNAi screens in *Drosophila* cell lines have provided insight into genes regulating various biological processes (8-15). However, this approach has drawbacks that limit resolution, including off-target effects and incomplete loss-of-function due to RNAi, and the high cost of reagent multiplicity and replication due to the arrayed format. Pooled CRISPR may address the major drawbacks: CRISPR generates complete loss-of-function alleles and causes fewer off-target effects on average (16, 17), and the pooled format allows greater multiplicity and replication for unit cost. Approximately half of the genes in *Drosophila*, arguably the best characterized multicellular genetic model system, lack functional characterization (18), so the need to develop orthogonal screening approaches is clear.

Here, we introduce a new method to deliver pooled DNA libraries stably into invertebrate cell lines. We use this technology to conduct a genome-wide CRISPR screen in *Drosophila* cells and show that the method identifies 1235 genes essential for fitness, 303 of which are uncharacterized in *Drosophila*. Moreover, we show that the system can be used in combination with drug perturbation to identify genes that when knocked out buffer cells against the drug or act synergistically with it. The method should be amenable to adapting any pooled DNA library screening approach to *Drosophila* or other invertebrate cell lines, such as shRNA knockdown (2) or CRISPR activation/inhibition.

## Results

Pooled mammalian cell-line screens use lentiviral vectors to deliver highly complex libraries of DNA reagents. However, the use of lentiviral vectors in insect cells is extremely inefficient (unpublished observations) possibly due to toxicity (19). An important advantage of library delivery using lentiviral transduction is that each sequence integrates into a euchromatic site in the host chromosome where it is expressed and remains in the genome as a detectable barcode such that enrichment or de-enrichment of each sequence can be monitored by massively parallel (next-generation) sequencing. As an alternative strategy, we assessed the efficiency of phiC31 site-specific recombination mediated by plasmid transfection. Methods for recombination in cell lines were recently developed (20-22), but the quantitative efficiency of integration has not been reported. Efficiency is critical for pooled screening applications as it must reach a threshold above which generating and maintaining >1000X representation of a library of tens of thousands of elements becomes technically and cost-feasible (23). We note that a previous study attempted to use pooled transient transfection for barcoded delivery of a DNA library in *Drosophila* cells but failed to identify any essential genes using the system, most likely due to the lack of a mechanism for retention of the DNA reagents in the cells through extended passaging (24).

To test phiC31 integration efficiency in *Drosophila* cells, a *Drosophila* S2R+ cell-line derivative, PT5, harboring a mobilized MiMIC transposon containing *attP* sites flanking mCherry (21) was transfected along with a plasmid containing *attB* flanking a GFP-2A-Puro(res) cassette, which we termed ‘pLib6.4‘, along with a phiC31 helper plasmid (Figure 1A). The population was then passaged for two months to dilute unintegrated DNA. Importantly, no selection reagent was added during these passages in order to monitor the efficiency of integration rather than the added efficiencies of integration and selection. Integration efficiency, inferred through flow cytometry (Figure 1B), suggested that phiC31-mediated cassette exchange occurred in ∼20% of the cells (even without accounting for incomplete transfection), 123-fold more than the background illegitimate recombination rate observed without phiC31 (Figure 1B). Interestingly, pLib6.4, which contains separated *attB* sites flanking the DNA sequence to be integrated, allowed ∼40-fold greater integration efficiency than traditional *Drosophila* attB40 vectors such as pCa4B (25), in which the *attB* sites are adjacent and require integration of the entire plasmid (data not shown). Growth in puromycin enriched for integrants (Figure 1B).

**Figure 1.**
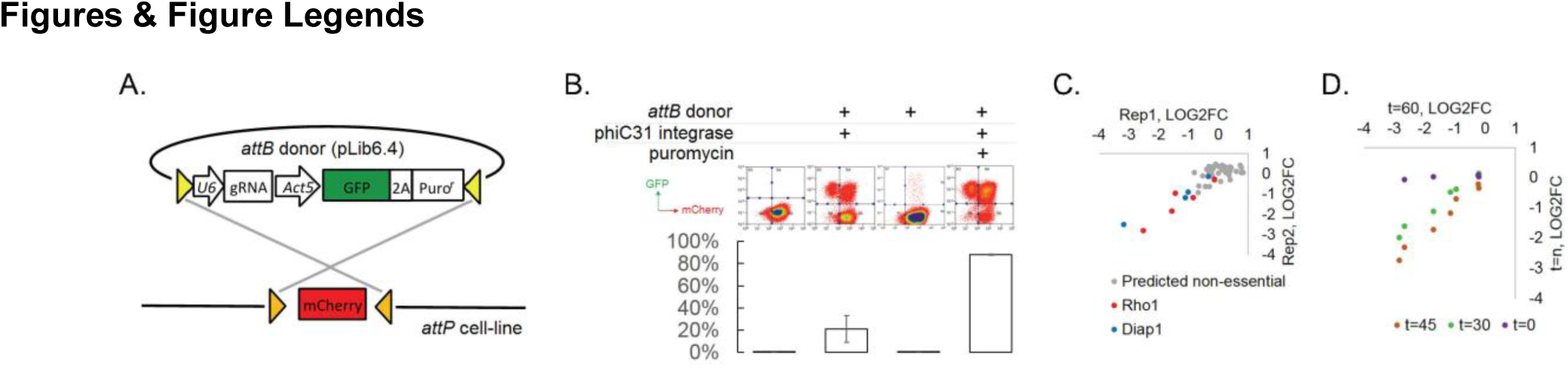
A novel method for introducing highly complex DNA libraries using phiC31 recombination. (A) phiC31 *attP*-*attB* recombination strategy. PT5 cells containing *attP* sites (gold) flanking mCherry were recombined with pLib6.4 plasmid library containing *attB* sites (yellow) flanking U6 sgRNA expression and GFP- 2A-Puro expression cassette. (B) Recombination efficiency measured by flow-cytometry. Indicated plasmids were transfected and grown with or without puromycin as indicated and passaged for 60 days. Graphs reflect total percentage of stable integrants (GFP+/total). N = 3; error bars reflect standard deviation. (C) Dropout of essential-gene targeted sgRNAs from a minipool of 31 sgRNAs. Two replicates of PT5 or PT5/Cas9 cells transfected with sgRNAs targeting *Rho1* (red) or *Diap1* (blue) along with sgRNAs targeting eight genes predicted to have non-essential functions (grey, see Suppl. Fig 1F) were passaged with puromycin for 60 days and sgRNA abundance was measured using next-generation sequencing. Graph shows log2 fold-change of each sgRNA in cells expressing Cas9 divided by sgRNAs in cells not expressing Cas9. (D) Optimizing passage time for dropout measurements. sgRNA abundance was detected from cells transfected as in (C) but 0 day, 30 days, or 45 days, and log2 fold-changes were compared to those at 60 days.

We next adapted the platform to CRISPR-Cas9 knockout screening. First, we generated PT5 S2R+ cells stably expressing metallothionein-driven SpCas9 (’PT5/Cas9‘). In combination with an sgRNA targeting the nonessential gene *Dredd*, PT5/Cas9 cells without induction were capable of editing nearly 50% of *Dredd* alleles, which was higher than that achieved by repeated rounds of transient transfection with a SpCas9 expression plasmid and not improved by copper induction (Suppl. Fig 1A,B). In a next test, we transfected cells with a pool of sgRNAs in pLib6.4 and monitored sgRNA abundance following passaging. After 60 days (roughly 60 cell doublings), the pools were significantly depleted of some sgRNAs targeting the essential genes *Rho1* and *Diap1* relative to those targeting genes predicted to have non-essential functions (Figure 1C). Monitoring *Rho1* or *Diap1*-targeted sgRNAs in the screen pools 30, 45, or 60 days post-transfection showed that 45 days of passaging is optimal (Fig 1D). To determine whether transfection multiplicity at the time of Cas9 expression affects essential gene-targeted sgRNA dropout efficiency, we developed an inducible Cas9 expression system in PT5 cells using intein-Cas9 (26) coupled with inducible expression in order to withhold Cas9 activity until sgRNAs have integrated (Suppl. Fig 1C-F). Surprisingly, a comparison of dropout efficiencies between the inducible and constitutive Cas9 platforms showed more selective reduction of *Rho1* or *Diap1* sgRNAs with constitutive rather than inducible Cas9, most likely due to the lower overall editing efficiency from intein-Cas9 as previously reported (27) (Suppl. Fig 1F). From these results, we conclude that the use of phiC31 integration into the PT5/Cas9 cells is suitable for scalable genome-wide perturbation screening using a pooled sgRNA library in *Drosophila* cells.

To construct a genome-wide sgRNA knockout library for *Drosophila*, we pre-computed sgRNAs for the first half of the coding region of all genes (26), applied efficiency and frame-shift filters previously shown to correlate with reagent success (15, 28), and ranked the remaining designs based on the uniqueness of the seed region sequence (12-bp downstream of the PAM) and the number of potential off-target sites. The top ranked 6-8 sgRNAs per gene (85,558 in total) were chosen and synthesized together on a microarray chip with non-targeting and intergenic negative controls, harvested by PCR, and cloned into pLib6.4 using published methods (29). We used a barcode strategy to separate the full library into three focused library pools so that we could perform focused or genome-wide screens. Each focused subset of the library targets a unique set of experimental genes as well as a common set of controls (Suppl. Fig 2A).

**Figure 2.**
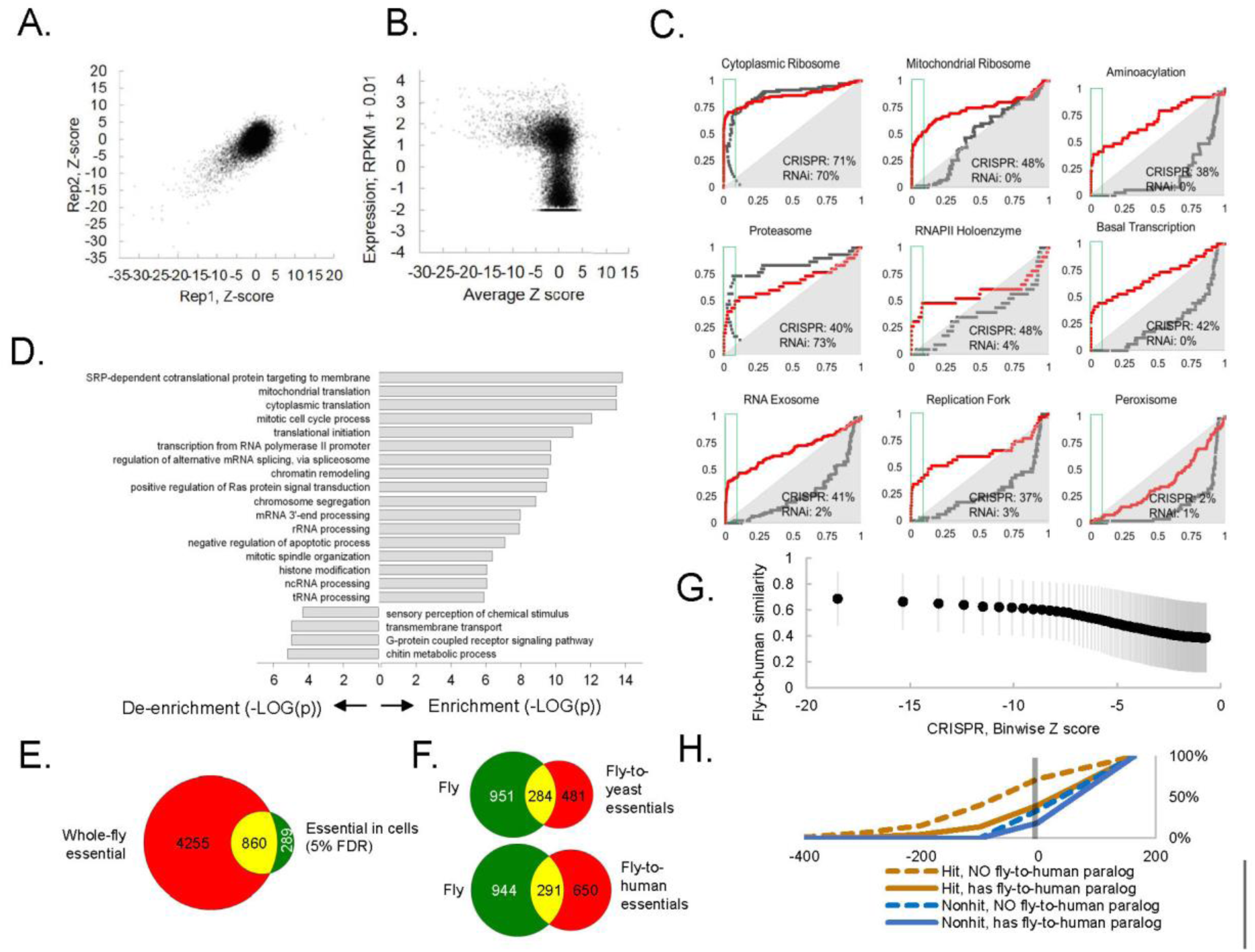
CRISPR/Cas9 knockout dropout screening in a *Drosophila* cell-line. (A) Dropout efficiency for each sequence of a library of ∼87,000 sgRNAs was measured using next-generation sequencing (see Materials and Methods) and log2 fold-changes were aggregated into a single Z-score using the maximum likelihood estimate (MLE) computational approach for each of 13,928 *Drosophila* genes in two independent, sequential replicates and plotted. (B) Z-scores for computed replicate averages (Supplementary Table 1) were plotted against RNAseq expression value (LOG10(RPKM + 0.010)) from the S2R+ cell-line (MODEncode). (C) Receiver operating characteristic (ROC) curves using mRNA-expression-based FDR (Suppl. Fig. 3A) versus rate of discovery of components of selected essential eukaryotic complex (55). Curves compare CRISPR knockout screen (this study) with reanalyzed genome-wide RNAi (8) (Suppl Fig. 2A). (D) Top enriched gene ontology (GO) terms for screen hits at 5% FDR compared top enriched GO terms for non-hits. (E) Overlap between cell-line CRISPR hits at 5% FDR and all “lethal” Flybase entries after subtracting entries with no allele information. (F) Overlap between CRISPR hits at 5% FDR and fly-to-yeast or fly-to-human fitness genes (see Materials and Methods). (G) Protein similarity correlation with Z score. A rank-based binning function was applied to CRISPR or RNAi data, where every 100 genes was binned, and each binwize average Z-score plotted against each binwize protein similarity value. Error bars reflect standard deviation. (H) Effect of fly-to- human paralogs on hit-calling in human CRISPR screens. Cumulative average of gene fitness essentiality (negative Bayes Factor) for high-resolution human cell-line CRISPR screen (7, 56) examining indicated genesets: those with paralogs are dashed; orthologs of fly fitness genes are brown; orthologs of non-hits are blue.

To identify fitness genes under basal growth conditions, we transfected cells with pLib6.4 containing the library of sgRNAs along with phiC31 helper plasmid and passaged the cells every 5 days for 45 days (Fig. 2A). We transfected ∼1,500 cells per sgRNA and carried >1,500 cells per sgRNA per passage to maintain the diversity of original library. To determine reagent correlation, the three separated sgRNA pools were each transfected and cells passaged independently and common controls were compared (Suppl. Fig. 2A; Supplementary Table 1). Comparing sgRNAs prior to transfection versus 45 days after transfection yielded high reagent correlations between the same sgRNAs as demonstrated for *Rho1* or non-targeting guides, for which different sequences produced a highly reproducible range of different dropout efficiencies (Suppl. Fig. 2B; Supplementary Table 1).

We used MAGeCK (30) maximum likelihood estimation (MLE) to compute fitness Z-scores for each gene from the log2 fold-changes of individual sgRNAs, which yielded good gene-level correlation between sequential biological replicates (Figure 2B). Because all true fitness genes must be expressed as mRNA, fitness scores were compared with gene expression from S2R+ cells (31) as an orthogonal validation of the CRISPR results (Figure 2C). Fitness genes were highly enriched for genes with any expression level of RPKM > 1 (Figure 2C; Suppl. Fig 3B). By ranking genes by fitness score, we calculated a false-discovery rate (FDR, defined as cumulative distribution of false-positive/[true positive + false positive]) as a function of rank and set a cutoff at 5%, identifying 1235 fitness genes (Suppl. Fig 3A).

**Figure 3.**
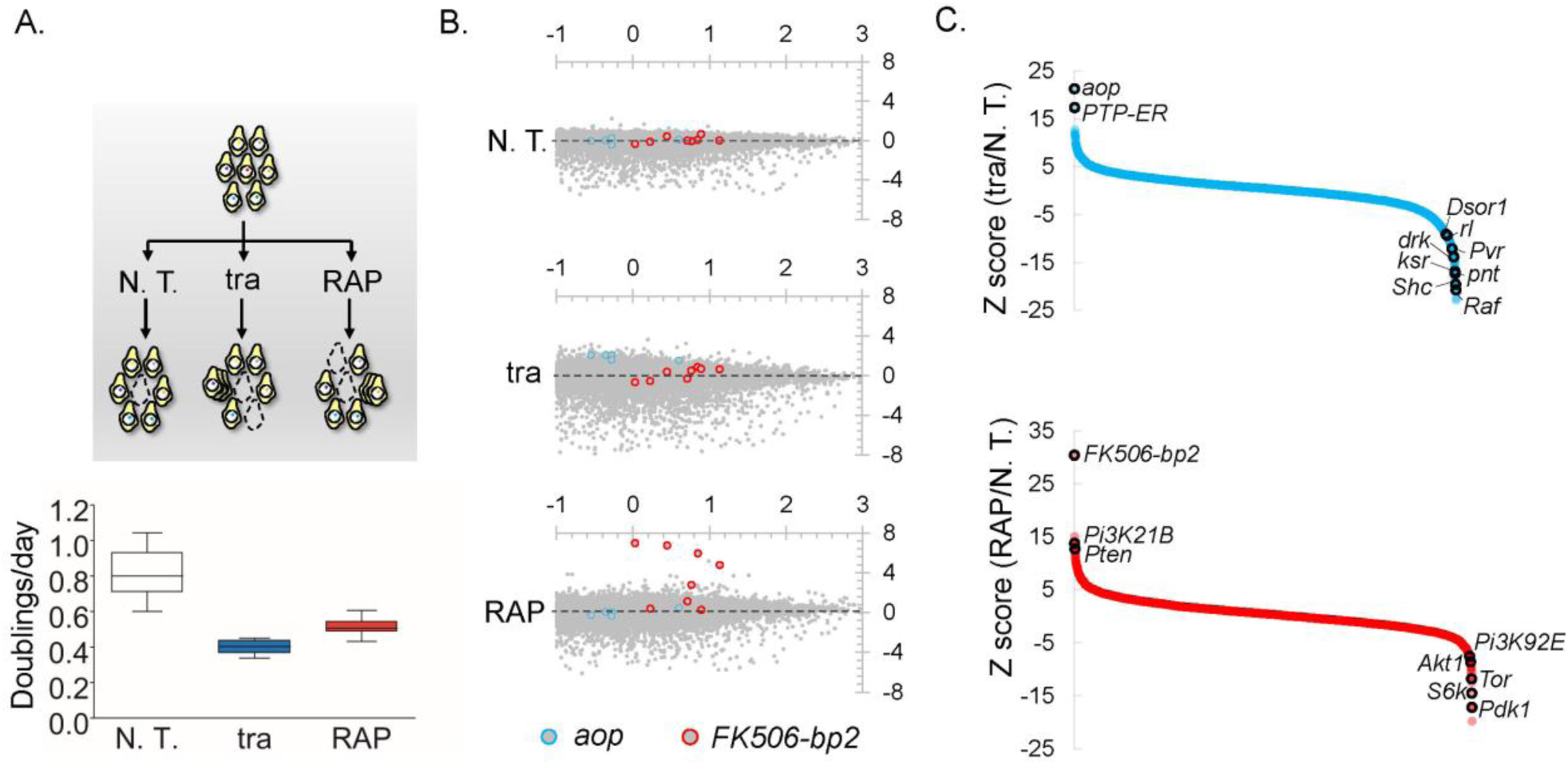
Screens to identify genes regulating cell growth and proliferation. (A) Schematic of experiment: pathway-specific perturbations to identify context-specific gene essentiality using *Drosophila* CRISPR screens. Trametinib (“tra”) suppresses the Ras/ERK/ETS pathway; rapamycin (“RAP”) suppresses the PI3K/mTOR pathway; while dropout screens conducted with no additional treatment (N.T.) serves as a control. Estimates of doubling per day obtained during periodic counting of cell pools to verify drug toxicity. (B) Plot of log2(initial distribution) versus log2(fold-change) for indicated sgRNAs in each screen. Pathway-specific resistance for sgRNAs targeting *aop*, a known suppressor of the Ras/ERK/ETS pathway in *Drosophila* (45) or *FK506-bp2*, the putative cellular co-factor for rapamycin (46). (C) Computed maximum likelihood estimate (MLE) Z score based on sgRNA fold-change data comparing drug treatment condition with no treatment control. Expected intra-pathway negative or positive regulators are highlighted (see Supplementary Table 3 for complete hit list and raw data).

The current state-of-the-art for *Drosophila* cell-based screens is arrayed RNAi (1). We re-analyzed a genome-wide RNAi viability screen in S2R+ cells (8) and removed double-stranded RNA designs with predicted off-targets, which significantly lowered the FDR of the RNAi screen (see Materials and Methods). Nevertheless, compared with pooled CRISPR screen results, the re-analyzed RNAi-based fitness gene set still had a far higher FDR, allowing the discovery of only 145 fitness genes at an FDR of 5% or incurring 44% FDR with a fitness gene set of 1235 genes (Suppl. Fig. 3A,B). Among genes with an RPKM > 1, RNAi was more likely to identify highly expressed genes, whereas this bias was less prominent in the CRISPR screen (Suppl. Fig. 3B,C). We used receiver-operating characteristic (ROC) curves to determine screen sensitivity for conserved, large macromolecular complexes. Whereas CRISPR and RNAi were both able to detect cytoplasmic ribosomal or proteasomal genes, CRISPR was better able than RNAi to identify significant fractions of the mitochondrial ribosome, aminoacyl-tRNA ligases, RNA polymerase II, basal transcription factors, RNA exosome, or the replication fork at low FDR (Figure 2D). As a control, neither method detected peroxisome genes, consistent with the observation that cell lines lacking peroxisomes grow normally (32) (data not shown) (Figure 2D).Interestingly, a human CRISPR-RNAi screen comparison also found a similar inability of RNAi to detect genes of the mitochondrial ribosome whereas it exceeded CRISPR sensitivity in detecting genes of the cytoplasmic ribosome, possibly due to differences in mRNA stability (7).

As an alternative method to demonstrate screen reliability, we performed gene ontology enrichment on CRISPR screen hits at 5% FDR. The gene set is enriched for essential processes such as translation, transcription, splicing, etc. and de-enriched for processes such as chitin metabolism that are necessary in organisms but not cells (Figure 2E). Finally, at 5% FDR, ∼17% of cell-essential genes overlapped with whole-fly essential genes, which were defined as genes with lethal alleles annotated in FlyBase (Figure 2H). This overlap is in a similar range as knockout mouse viability compared with mouse cell-line fitness essentiality (where the overlap is ∼27%) (33), and may represent different genetic requirements of whole animals versus cell lines or of germline versus somatic cells (34).

Copy number has been shown to correlate with the CRISPR viability score in some mammalian CRISPR screens, presumably due to induction of greater DNA damage foci for copy-number-amplified (CNA) loci, creating spurious fitness calls (35). By contrast, we find that CRISPR in *Drosophila* shows no detectable bias towards CNA genes for 97% of the genome (Suppl. Fig. 3D). Interestingly, genes with extreme CNA of 8 or more copies (3% of genes) exhibited a bias of ∼1.8 fold in the CRISPR screen, but the magnitude of this bias was similar to that seen in an RNAi screen for the same genes (Suppl. Fig 3D), suggesting that CNA genes in *Drosophila* cells are enriched for fitness essentiality. Finally, copy-number correction had no detectable effect on FDR, true-positive rate, or enrichment of major macromolecular complexes (not shown). The analysis shows that *Drosophila* CRISPR screens do not have detectable copy-number bias.

A technical comparison of CRISPR screens in humans and fly cells indicated that the method performs similarly in both systems. First, the fly screen had similar sensitivity (true-positive rate at 5% FDR) to genome-wide screens in human cell-lines performed with similar reagent number (GeCKO v2 screens) when compared against top RNAi hits (36) (Supp. Fig. 3E). By comparing sgRNAs that dropped out versus those that failed to dropout for known essential genes, we were able to compute a probability-based position matrix for optimal sgRNA design in *Drosophila* (Suppl Fig. 2E). The position matrix shows complementary nucleotide biases at 15/21 positions and is broadly consistent with human CRISPR screens (37) (Suppl. Fig. 3F). Specifically, out of the first position of the PAM sequence and the 8 bp seed region before the PAM sequence, 5 of 9 positions are fully consistent while 2 of 9 position are partially consistent with the Doench score (37) (Suppl. Figure 3F). The analyses support an underlying mechanistic unity of targeting by Cas9 and NHEJ repair between human and fly cells.

A high-resolution fitness map in fly cells now allows us to compare fitness genes among species. The fly essential gene set partially but significantly overlaps with characterized essential gene sets from yeast and human cells (Figure 2F). Moreover, fitness essentiality correlates with protein conservation in humans (Figure 2G). Despite significant overlap between fly and human cell-line fitness genes and conservation of hits in both screens, many fly fitness genes map to human genes that were not identified in human CRISPR screens. We hypothesized that fly CRISPR screens identify essential genes in *Drosophila* cells but not in human cells due to paralog redundancy. To test this hypothesis, we inferred putative paralog relationships for moderately or highly conserved orthologs using the DRSC Integrative Ortholog Prediction Tool (DIOPT) (38). Note that we only considered genes that were expressed as mRNA in the human cell-line (Figure 2H; Supp. Figure 4). Using this framework, 35% of conserved human genes contain a fly-to-human paralog, and 407 fly fitness genes (at 5% FDR) are the orthologs of human genes containing at least one paralog. Interestingly, in human cell-line CRISPR screens, there is significant bias against detecting genes that are conserved in flies and have a paralog in the human genome, and this bias was dramatically elevated for the human orthologs of fly fitness genes (Figure 2H; Supp. Figure 4). Thus, we propose that redundancy among closely related proteins in cell-line screens reduces the sensitivity of human CRISPR screens, suggesting that lower complexity *Drosophila* cells can identify pairs of paralogous genes required for human cell fitness.

Although the *Drosophila* fitness genes we identified are enriched for characterized phenotypes and publication count relative to all genes (data not shown), phenotypes have yet to be described for 303 of them. Among these 303 genes, 251 have human orthologs (Supplementary Table 2). Thus, further studies of these conserved genes are likely to provide new insights into conserved, cell essential processes not yet studied in flies. Also of interest are fly-specific fitness genes, as they present a paradox and may reveal novel species-specific biology or overlooked structural/functional analogs without sequence orthology and may have potential as targets for new insecticides. At 5% FDR, we obtained 62 fitness genes with no sequence similarity outside of flies using DIOPT, and phenotype information exists regarding 25 of these (Supplemental Table 2). In confirmation of our methods, these included known cell-essential divergent genes such as *ver* and *HipHop*, which encode components of the telomerin complex, the putative functional analog of mammalian telomerase (39, 40), and *Kmn1, Kmn2* and *Spc105R*, whose gene products may be structural anologs of Ndc80, and Mis12 complex components that interact with centromeric DNA, a proposed driver of speciation (41-43), as well as several chromatin-interacting proteins (Supplemental Table 2). The characterization of the remaining conserved and non-conserved genes is likely to bring new insights into essential gene functions.

We next asked whether our CRISPR screening platform could be used to identify genes acting in signaling pathways that regulate cell growth and proliferation (44). To do this, we used two focused libraries targeting a total of 3,977 genes in positive selection screens for growth in the presence of trametinib (tra), an inhibitor of the Ras/ERK/ETS pathway, or rapamycin (RAP), an inhibitor of the mTOR/PI3K pathway, with the aim of uncovering known and novel compensatory mechanisms or synergizing loss-of-function mutations (Fig 3A). Each drug was applied to screen pools at a moderate dosage that still allowed cells to double but at approximately half the normal rate and then pools re-sequenced. The effect of each drug was carefully monitored by periodically counting cells during the screen to confirm the effect on cell doubling rate (Fig 3A). As an illustration of screen specificity, sgRNAs against *aop*, a transcriptional repressor of the Ras/ERK/ETS pathway (45), or the putative intracellular co-factor for rapamycin, *FK506-bp2* (46), conferred resistance specifically in the context of tra or RAP, respectively (Figure 3B). We focused on other such genes for which multiple sgRNAs provide a survival benefit or synergistic lethality in the context of drug treatment using maximum likelihood estimate (30) (MLE) (Figure 3C; Supplemental Table 3). The analysis identified known negative regulators of each pathway among the set of genes that conferred resistance when knocked out, and identified several pathway positive regulators as synthetic lethal (Figure 3C). The data additionally revealed cross-pathway synergy and several novel candidate pathway regulators (Supplementary Table 3) and demonstrates that CRISPR screening in *Drosophila* cells can reveal context-specific compensatory mechanisms or essentiality relevant to major signaling pathways.

## Discussion

CRISPR screening technologies have illuminated the functions of uncharacterized genes and provided a straightforward, cost-effective pipeline to evaluate gene function in different contexts (47). However, the benefits of this approach have been inaccessible for *Drosophila* and other insects because the delivery of highly multiplexed DNA libraries has thus far required lentiviral transduction, a process that fails to produce transformed cells (unpublished and (1)). Here, we show that multiplexed DNA delivery by transfection followed by site-specific recombination is an effective alternative strategy. We use this technique to deliver a genome-wide library of sgRNA expression cassettes and perform CRISPR knockout fitness screens in *Drosophila* S2R+ cells, identifying 1235 fitness genes at 5% FDR. Moreover, our *Drosophila* CRIPSR knockout system identified mutants conferring drug resistance or synergy, and should thus be suitable for many future context-specific fitness experiments examining gene-drug/nutrient or gene-gene interactions.

The introduction of systematic knockdown and knockout approaches have greatly reduced false-positive assertions in human cell-line loss-of-function studies, but equally important is knowing what genes are missed by these approaches and providing alternative screening strategies that can target them. Of critical importance are paralogous genes with redundant functions, i.e., the presence of one paralog buffers against the loss of another (18). Although relevant to human disease (48), these genes are “invisible” in singe-gene screens and dramatically reduce search space in gene-by-gene screens (18). When we compared human and fly CRISPR screens, human CRISPR screens were less able to detect the fly orthologs of fitness essential genes when those genes had paralogs relative to the fly genome. This supports the use of parallel *Drosophila* and human screens as one approach to offset genetic redundancy.

In addition to CRISPR knockout, the strategy we report can be used with numerous other high-throughput screening modalities which were previously not possible in *Drosophila*, including CRISPR activation, CRIPSR inhibition, CRISPR base-editing, shRNA knockdown, cDNA overexpression, perturb-SEQ, and combinatorial approaches for multigene suppression/activation(47, 49, 50). Finally, the methods and constructs we employ are likely to be directly transferable to the large number of existing cultured cell-lines derived from other insects such as mosquitos, where they could be used to characterize viral propagation mechanisms in the vectors of human pathogens such as Dengue or zika.

## Materials and Methods

### Vectors and cell lines

pBS130, phiC31 integrase under control of the HSP70 promoter, was obtained from Addgene. Transient Cas9 expression used pl018 (15) containing *Drosophila*-optimized Cas9 under the strong Actin promoter. Cas9 from pl018 followed by the BGH terminator from pcDNA3.1 (Invitrogen) were cloned into the SpeI/NotI site of pMK33 (51) to generate pMK33/Cas9. Human codon-optimized intein-Cas9_S219- 3XFLAG (52), a kind gift of D. Lui, was amplified by PCR and cloned into pMK33 to generate pMK33/intein- Cas9_S219-3XFLAG. pLib6.4 was generated by using standard cloning methods as follows. First, the *Drosophila* U6:2 and Act5C promoter cassette from pl018 (15) was amplified by PCR using primers containing the 5’ attB40 site and inserted into pUC19 using Gibson assembly. Next, GFP-T2A-Puro from pAc-STABLE2 (53) was amplified with primers containing the 3’ attB40 and introduced by ligation into an engineered site in the resulting vector and verified by Sanger sequencing. For minipool experiments in Figure 1, sgRNAs were cloned individually using standard methods into the BpiI/BbsI site of pLib6.4 and verified by Sanger sequencing and then mixed. Sequencing reactions were carried out with an ABI3730xl DNA analyzer at the DNA Resource Core of Dana-Farber/Harvard Cancer Center (funded in part by NCI Cancer Center support grant 2P30CA006516-48). S2R+ derivative PT5 (NPT005) was obtained from the *Drosophila* RNAi Screening Center (21) and grown in Schneider’s medium (Thermo Fisher Scientific) containing 10% heat-inactivated FBS. PT5 cells were transfected with pMK33/Cas9 or pMK33/intein-Cas9_S219-3XFLAG and selected over a period of two months in 200 ng/uL Hygromycin B (Calbiochem). The resulting PT5/Cas9 cells were maintained in Hygromycin until CRISPR library transfection.

### Library design and construction

In order to allow focused sublibrary screening as in Figure 3, sgRNA library was encoded in three separable sublibraries with common controls (Suppl. Figure 2A). Library synthesis was performed by CustomArray using published methods (5, 29). Briefly, CRISPR sgRNAs were encoded within a 110-nt single-stranded DNA oligo containing unique library-specific barcode sequences and flanked by BpiI/BbsI sites. 15-cycles of PCR using Phusion Polymerase (New England Biolabs) were used to amplify each sub-library, and then BpiI (Fermentas) was added to the amplicons. A non-denaturing 20% TBE gel (Thermo) was used to purify the resulting 23-mer fragment and eluted overnight using the crush-soak method. The concentration of ligatable fragments in each preparation was empirically optimized by test ligations in pLib6.4. Optimized concentrations were used in ligations with BpiI-digested pLib6.4 and then electroporated into Ecloni 10GF’ electrocompetent cells (Lucigen) at a yield of 20-50 times diversity for each library, generating dense colonies on ∼60 150-mm LB-carbenicillin plates, which were grown for 18 hours at 37°C and harvested by scraping into LB medium followed by mixing suspended colonies by extensive vortexing. Glycerol was added to 50% and the libraries were flash frozen in 1 mL aliquots. Each sublibrary was prepared by pooled minipreps and eluted in buffer EB (Qiagen).

### Transfection, pool maintenance, and drug addition

pBS130 and pLib6.4 containing CRISPR sublibraries were mixed and co-transfected into PT5/Cas9 cells using Effectene with the manufacturer’s recommended protocol (Qiagen). For each transfection, cells were first grown until just confluent on T75 flasks for 2-3 days. Doubling was monitored at this passage and determined to be approximately 1/day. Then cells were removed by forceful tapping and replated at 3 X 10^6^ per well into 6-well dishes. The number of dishes required reflected the library size and accounted for incomplete transfection efficiency to give at least 1500 cells/sgRNA (for reference, a full-genome screen required all wells of eight 6-well dishes). t=0 used plasmid read count values. Transfection efficiency and integration efficiency were monitored periodically after transfection for the first twelve days using flow cytometry (BD LSRII). Flow cytometry was performed at the Harvard Medical School Department of Immunology Flow Core. Each 1.5 wells of cells was transferred to one 10 cm dish and passaged in the presence of 5 ug/mL puromycin and cells were allowed to become confluent over 4-6 days.Next, the pool was contracted by two-fold, and 2 x 10^7^ cells from each of two plates was combined into a single 10 cm dish. 2 x 10^7^ cells were passaged every 5 days until downstream processing at the indicated time.These steps during early phases of selection ensured no loss of diversity due to variable transfection efficiency. To determine partially inhibitory dose of trametinib (Selleck) or rapamycin (Tocris), the effect of varying doses of each drug was first measured in a 4-day treatment to PT5/Cas9 cells by Cell Titer Glo assay (Promega), using manufacturer’s recommended protocol (not shown). From this data, 50 nM trametinib or 2 nM rapamycin was chosen as partially perturbative concentrations for focused library screens (Figure 3). To verify that drug treatment decreased doubling rate during the screen, cell counts were performed periodically (Figure 3B).

### Library sequencing and CRISPR screen data analysis

Genomic DNA was prepared using the Zymo miniprep kit. Assuming DNA content in S2R+ cells was 4N, each cell contains ∼0.6 pg of DNA. To maintain diversity, all PCRs were conducted from ∼5,000 cells per sgRNA per condition. Library amplification was performed using a two-step procedure. First, in-line barcoded inside primers (PCR1F x PCR1R) were used to amplify the library in 23 cycles such that the resulting amplicon had the following sequence: 1/2Read1- (N)_n_ANNEALINGSEQUENCE-sgRNA-tracrRNA. Primer sequences conformed to: 5‘-CTTTCCCTACACGACGCTCTTCCGATCT(N)_n_ (B)_6_ gttttcctcaatacttcGTTCg-3’ (where N=any nucleotide; n=variable number between 1-9; and B=barcode nucleotide) and 5‘- TTTGTGTTTTTAGAATATAGAATTGCATGCTGggtacctc-3‘. Next, common outside primers were used to amplify these amplicons with an additional 11 cycles such that the resulting amplicons had the following sequence: P5-Read1-(N)_n_-ANNEALINGSEQUENCE-sgRNA-tracrRNA-P7. Sequences were: 5‘- AATGATACGGCGACCACCGAgatctACACTCTTTCCCTACACGACGCTCTTCCGATCT -3’ and 5‘- ATCTCGTATGCCGTCTTCTGCTTGTTTGTGTTTTTAGAATATAGAATTGCATGCTGggtacctc-3‘. Following second amplification, amplicons were gel purified and concentration was determined using Qubit dsDNA HS Assay Kit (Thermo). Amplicons were pooled according to concentration of sgRNAs per unit volume. Pooled barcoded amplicons were subjected to sequencing using the NextSeq500 1 X 75 SE platform (Illumina).Sequencing was performed at the CCIB DNA Core Facility at Massachusetts General Hospital (Cambridge, MA) or the Harvard Medical School Biopolymers Facility (Boston, MA). De-multiplexing of the library was performed using Tagdust (54) and all downstream analysis was performed within MAGeCK (30) with the following experiment-specific changes: 1) Read count files were stripped of sgRNAs below the 10th percentile lowest initial readcounts for each sublibrary before processing in MAGeCK MLE we performed 1000 iterations for all Z-score assignments. For drug selection experiments, several context-nonspecific sgRNAs providing survival benefit to cells under normal growth conditions were removed due to wide variation of these sgRNAs upon selection.

### Bioinformatics analysis

For ROC curves, major eukaryotic essential complex components for *Drosophila* were from KEGG (http://www.genome.jp/). For gene ontology enrichment analysis, annotation file for *Drosophila* genes was retrieved from NCBI (ftp://ftp.ncbi.nlm.nih.gov/gene/DATA/gene2go.gz). Hypergeometric analysis was done to calculate the enrichment P value using in-house program written in JAVA.Ortholog assignment as well as fly-to-human protein similarity score were obtained using DIOPT v 6.0.2 using “moderate” or “high” confidence calls (38). RNAseq analysis in this paper used the webtools DGET (http://www.flyrnai.org/tools/dget/web) or CellExpress (http://www.flyrnai.org/cellexpress), which used RNAseq expression data from modENCODE (31).

### Data Availability

FASTQ files and readcount files for CRISPR analysis compatible with MAGeCK are available upon request. pMK33/Cas9, pMK33/intein-Cas9_S219-3XFLAG, and pLib6.4. are available through Harvard PlasmID Database (http://plasmid.med.harvard.edu). The three CRISPR sublibraries used in this study are available through DRSC/TRiP Functional Genomics Resources (https://fgr.hms.harvard.edu/).

## Acknowledgments

We thank S. E. Mohr and B. E. Housden, and B. Ewen-Campen for useful discussions. This work was supported by the LAM foundation (LAM0122P01-17) and the Department of Defense (W81XWH-16-1-0127).R. V. received support from Harvard Medical School Department of Genetics NIH Training Grant(5T32GM007748-39). Y. H. is supported in part by NIH grant NIGMS R01 GM067761. N.P. is a Howard Hughes Medical Institute investigator.

